# Navigating the Unstructured by Evaluating AlphaFold’s Efficacy in Predicting Missing Residues and Structural Disorder in Proteins

**DOI:** 10.1101/2024.11.03.621778

**Authors:** Sen Zheng

## Abstract

This study explored the difference between predicted structure confidence and disorder detection in protein, focusing on regions with undefined structures detected as missing segments in X-ray crystallography and Cryo-EM data. Recognizing the importance of these ‘unstructured’ regions for protein functionality, we examined the alignment of numerous protein sequences with their resolved or not structures. The research utilized a comprehensive PDB dataset, classifying residues into ‘modeled’, ‘hard missing’ and ‘soft missing’ based on their visibility in structural data. By analysis, key features were firstly determined, including confidence score pLDDT from AlphaFold2, an advanced AI-based tool, and IUPred, a conventional disorder prediction method. Our analysis reveals that "hard missing" residues often reside in low-confidence regions, but are not exclusively associated with disorder predictions. It was assessed how effectively individual key features can distinguish between structured and unstructured data, as well as the potential benefits of combining these features for advanced machine learning applications. This approach aims to uncover varying correlations across different experimental methodologies in the latest structural data. By analyzing the relationships between predictions and experimental structures, we can more effectively identify structural targets within proteins, guiding experimental designs toward areas of potential functional significance, whether they exhibit high stability or crucial unstructured regions.

## Introduction

Proteins are fundamental to life, and understanding their three-dimensional (3D) structures is crucial for elucidating their functions. Structural biology has made significant strides and underscoring the central dogma that a protein’s function is intricately linked to its 3D structure [1, 2]. These advancements are reflected in resources such as the Protein Data Bank (PDB), which archives extensive structural data essential for research [3].

Three primary experimental approaches are foundational in determining protein structures: X-ray crystallography (X-ray), Nuclear Magnetic Resonance (NMR) spectroscopy, and cryogenic electron microscopy (cryo-EM). X-ray crystallography requires proteins to be crystalline, which can be challenging for flexible regions and complex assemblies [4]. NMR spectroscopy offers insights into molecular flexibility but is primarily limited to smaller proteins [5]. Cryo-EM has revolutionized the field by enabling 3D reconstructions in near native states without the need for crystallization, positioning it as a prime candidate for atomic-level resolution in the future [6]. As a developing tool, cryo-EM includes modalities like single particle analysis (SPA) and tomography (Tomo), facilitating diverse structural exploration strategies [7, 8]. Structural biology now offers a sophisticated array of techniques that allow researchers to tailor their approaches based on protein characteristics such as size, assembly complexity and function, as well as specific experimental objectives and conditions.

However, many proteins have regions that are not strictly structured, known as disordered regions [9]. This partially or fully “unstructured” can arise for various reasons, such as inconsistencies in X-ray scatter patterns [10, 11] or the challenges of reconstructing precise models from fragmented density maps in Cryo-EM [12, 13], due to the intrinsic dynamic nature [14]. Analysis and indirect structural studies have revealed that these ambiguous areas are crucial for many protein functions [15, 16]. These proteins, marked by their variability and absence of a defined structure, are called intrinsically disordered proteins (IDPs) and regions (IDPRs) [17]. Their flexibility allows them to adopt multiple shapes, enhancing their versatility and effectiveness. Despite their lack of rigidity, they are flexible, biologically active, and essential for various cellular functions.

Classifying and predicting protein disorders is therefore crucial, given the limited and time expansive experimentally determined structures and the vast sequence data in expanding UniProt [18]. Traditional disorder predictors formally used bioinformatics and sequence analysis to help understand protein flexibility where integrate structures were scarce [19]. Among these tools, IUPred assesses interaction energies within amino acid chains, and havs proven effective for identifying disordered proteins and regions unlikely to form stable interactions [20, 21]. IUPred provides a complement to sequence-based methods, maintaining its status as a reliable comparator among disorder predictors [22].

In 2022, the field of protein structure prediction experienced a significant transformation with the introduction of AlphaFold2, a cutting-edge approach powered by deep learning [23]. Impressively, AlphaFold2 not only shines in pinpointing protein structures but also offers valuable insights into identifying IDPs and IDPRs, using its predicted Local Distance Difference Test (pLDDT) scores [24, 25]. This innovative method has sparked considerable interest for its potential in accurately spotting disordered areas within proteins [26, 27].

The advent of AlphaFold2 has revolutionized the field of structural biology with its advanced protein folding predictions. Despite its success, a comprehensive comparison between its ability to detect disordered protein regions and the performance of established experimental techniques like X-ray and Cryo-EM remains limited. These methods differ in pinpointing protein disorder, making it critical to assess AlphaFold2’s proficiency in this area. Our study addresses this issue by examining AlphaFold2’s performance in identifying disordered regions across structural experiments. We compare it with traditional tools such as IUPred, considering data from both past studies and recent protein discoveries since AlphaFold2’s inception.

To improve the understanding of protein structure representation, we developed a comprehensive framework evaluating computational prediction and validated on experimental results. Starting by curating a dataset of hard-to-structure protein structures and categorizing residues into "modeled", "hard missing", and "soft missing" classes, we aimed to elucidate the intricate relationships between predictive modeling, disorder, and structural determination. Our investigation revealed that "hard missing" residues often reside in low-confidence regions, but are not exclusively associated with disorder predictions. Leveraging pLDDT scores and machine learning, we also identified key features distinguishing "modeled" from "hard missing" residues, with varying correlations across experimental methodologies, notably SPA from Cryo-EM. Our findings demonstrate that complementary approaches can bridge gaps in structural prediction, particularly for dynamic and unstructured protein regions, informing the design of targeted structural experiments.

## Materials and Methods

### Dataset

Given the diverse protein functions and categorizations of disorder [10, 28], selecting a well-accepted definition to represent disorder that aligns with experimental and historical structures is crucial. To overcome and simplify the complexity and achieve our objectives, we adopted the basic definition of disorder as "missing coordinates" within the longest structural representations. This definition could provide a direct and unambiguous representation of structural disorder across structural experiments [28, 29].

We began by identifying the longest modeled protein entries and curated a dataset by compiling a list of protein entries with 100% sequence identity from the PDB as of 2024, retrieved via the wwPDB. By evaluating missing coordinates collected, we aimed to capture the fundamental meaning of disorder in structural experiments with focus on “hard-to-structure” regions over a broad and historical dataset.

Protein entries were consolidated and de-duplicated by grouping them under shared UniProt accession numbers, resulting in unique protein categorizations. These were further organized into three experimental method groups: X-ray crystallography, SPA and Tomo. Tomo group includes both tomography and sub-tomogram averaging methodologies.

The corresponding AlphaFold-predicted structures were downloaded from the AlphaFoldDB (www.alphafold.db). We obtained additional protein information from UniProt, including sequence in full-length, structural attributes, and secondary structures.

To ensure data integrity, we conducted a comprehensive review to align sequencing data among PDB, AlphaFoldDB, and UniProt, eliminating any discrepancies. This process finally culminated in a collection of 34,568, 11,072, and 416 unique protein entries for the X-ray, SPA, and Tomo datasets, respectively.

### Residue and Region Classifications

To determine which residues are fully or partially missing in datasets, we categorized residues obtained from PDB files into three groups based on the examining the presence of "Cα" carbon coordinate. Residue classification was as follows:

Modeled: Residues for which the "Cα" is present in all relevant entries for the same residue position.

Hard Missing: Residues for which the "Cα" is absent in all related entries.

Soft Missing: Residues for which the "Cα" is sometimes absent in the related entries.

In total, we identified 9,362,180/3,008,105/124,147 modeled residues, 546,152/875,223/ 32,636 hard missing residues, and 101,272/78,604/3,406 soft missing residues in X-ray/SPA/Tomo experiments, respectively. Given our focus on "hard-to-structure" residues, we prioritized entries with high residue counts, reflecting a higher prevalence of "hard" than "soft" missing components.

For region classification, we assessed the continuum of modeled, hard missing and soft missing residues to define corresponding regions. Both hard and soft missing regions were further categorized based on their length: regions with fewer or equal as 30 residues were classified as “short” disorder regions, while those containing more than 30 residues were classified as “long” disorder regions [30].

### Feature Extraction and Correlation Analysis

Protein features, including full-length sequences and secondary structure information, were extracted from their corresponding UniProt entries text. The pLDDT scores for protein sequences were obtained from structure files downloaded from AlphaFoldDB. IUPred scores were calculated using the locally installed IUPred3 software [21] with the "long method" applied to the respective protein sequences. In special and few cases where the length of a protein was less than 16 amino acids, the IUPred could not process, and these proteins were assigned a score of 0 at each residue.

To investigate the relationships between the extracted features and different residue groups, we counted the residues of each type, including modeled, hard missing, and soft missing residues. We then conducted Pearson correlation analysis on these counts against the counts of residues for each feature separately.

For a more detailed assessment, quadrant analysis were performed by categorizing the residues into four distinct groups based on their pLDDT and IUPred scores and plotted as instruction with some modification to suit our datasets [31]: Quadrant 1 (Q1), Residues with high pLDDT confidence (≥ 70) and low IUPred disorder (< 0.5); Quadrant 2 (Q2), Residues with high pLDDT confidence but high IUPred disorder ( ≥ 0.5); Quadrant 3 (Q3), Residues with low pLDDT confidence (< 70) and low IUPred disorder; Quadrant 4 (Q4), Residues with low pLDDT confidence and high IUPred disorder.

### Prediction Model Preparation and Training

All protein entries from X-ray, SPA, and Tomo groups were initially divided into two main sets. From dataset, protein entries containing any PDBs released before 2022 were designated as the training set, while those containing only PDBs released after 2022 were reserved for model validation.

For the basic pLDDT model, we followed general instructions [27, 29] and observations in this study, then make basic assumption that residues with a pLDDT score under 70 to be labeled “predicted” as "hard missing", otherwise “predicted” as "modeled". Similarly, in the basic IUPRED model, residues with IUPRED scores over 0.5 were “predicted” as "hard missing", otherwise as "modeled". The "soft missing" group was excluded from classification using the basic pLDDT and IUPRED models, as correlation was difficult to be detected found in this dataset.

To prepare for the training and further validation of LSTM model, a basic constructing progress was conducted with several modifications to meet our cases [32], the protein sequences along with their corresponding pLDDT and IUPRED scores were padded to a length of 2,500. The protein sequences were first tokenized into 21 dimensions, with an additional 2 dimensions corresponding to the pLDDT and IUPRED scores. Following the input layer, a hidden layer with 300 units was implemented, followed by a dense layer with soft-max activation. The target was defined as the classification of the true labels of residues into one of three categories: "modeled", "hard missing" or "soft missing" which were supplied to the final classification layer. The training optimizer employed was "adam", the loss function used was "categorical_crossentropy".

To balance the training dataset and prevent overfitting during this progress, all training sequences with fewer than 5 "hard missing" residues were excluded. Ultimately, two separate LSTM models were trained using the X-ray and SPA training datasets, respectively. Due to a scarcity of data, the Tomo LSTM model was not trained; instead, the trained SPA LSTM model was utilized for validation of Tomo group.

### Validation of Model Prediction

To validate the performance of residue group prediction, we assess the pLDDT, IUPred, and LSTM models using the validation dataset. The models are employed to predict residue groups, and we subsequently calculate the precision, recall, and F1 score for each model. This is accomplished by counting the true positive (TP), true negative (TN), false positive (FP), and false negative (FN) results from the predictions [32].

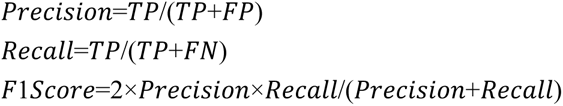

For residue group prediction, consider the example of the “modeled” group. The true positives and true negatives represent residues that are correctly classified as belonging to or not belonging to the “modeled” group, respectively. A false positive occurs when a residue not belonging to the “modeled” group is incorrectly classified as “modeled”. Conversely, a false negative arises when a residue belonging to the “modeled” group is misclassified as not “modeled”.

In a similar manner, for region group prediction, such as the “modeled” group region, true positives and true negatives refer to regions that are correctly predicted to overlap or not overlap with the “modeled” group by at least 50%. A false positive in this context is a region not belonging to the “modeled” group that is incorrectly predicted as “modeled”, while a false negative is a region of the “modeled” group that is predicted as not “modeled”.

### Analysis of Human known and unknown dataset

To evaluate the performance of our classification models on most valuable and well-studied data, we collected human protein sequences from UniProt, along with their pLDDT and IUPRED scores. Our objective was to identify high-confidence regions that could potentially be modeled but have not yet been characterized in this step. We categorized the collected entries into two primary classes: “Unsolved Sequences”: This class includes entries with recorded PDB structures. The sequence aligned referring to existing PDB entry and labeled as “solved sequence”. The remaining sequence that does not have corresponded PDB structural information are classified as “unsolved sequences.” This category contains a total of 8,099 entries that are containing partially unsolved in terms of structure.

“Unstructured Sequences”: This class encompasses entries with no associated PDB structures.

When a protein entry lacks any PDB representation, the entire sequence is classified as an “unstructured sequence”. As of March 2024, this category comprises 11,665 entries with no PDB representation.

Our analysis is to evaluate the potential “modeled” residues within “unstructured sequences” and “unsolved sequences”, and to find out distinguished high ratio of “modeled” ones. To predict the “modeled” residues, both basic pLDDT model and LSTM model were used. Finally, we plotted the predicted “modeled” residue rates based on the classification result of basic pLDDT model and the LSTM model (SPA based), for both the “unstructured sequences” and “unsolved sequences” of human proteins.

### Software

The main scripts used in this study, including data preparation and feature extraction, were written in Python. Pearson correlation coefficients were calculated using the Pandas package. LSTM models were trained using the TensorFlow and Keras packages. Visualization of the results, including violin and heatmap plots, was accomplished with Matplotlib and Seaborn.

## Results

### Distribution of Modeled and Missing Residues Across Experimental Method

To understand the distribution of missing residues and regions, we analyzed various structural determination methods results with regard to their unstructured segments, highlighting their difference in managing structural variability and disorder.

First, we assessed the frequency and length of "hard" and "soft" missing regions, which are defined by absent or partially present in structure within our datasets. In the X-ray dataset, 23% of entries were without "hard missing" regions and 82% lacked "soft missing" regions (Table 1, "no-missing" row). In contrast, SPA showed the lowest number of entries free of "hard missing" regions at 12%, while 90% were devoid of "soft missing" regions. Similarly, Tomo findings closely aligned with X-ray for "hard missing" regions at 25%, and recorded the highest absence of "soft missing" regions at 94%.

**Table 1.** Distribution of missing regions in various structural experiments.

|  |  | Xray |  | SPA |  | Tomo |  |
| --- | --- | --- | --- | --- | --- | --- | --- |
|  |  | Hard | Soft | Hard | Soft | Hard | Soft |
| **No-missing** |  | 23 | 82 | 12 | 90 | 25 | 94 |
| **Missing** | *Short* | 94 | 98 | 74 | 88 | 77 | 86 |
|  | *Long* | 6 | 2 | 26 | 12 | 23 | 14 |

Next, we categorized entries with "hard" or "soft" missing regions by length, dividing them into short ( ≤ 30 residues) or long (> 30 residues) sections. X-ray predominantly featured short "missing" regions, with 94% of "hard missing" and 98% of "soft missing" regions under 30 residues, indicative of limited tolerance for extensive disordered segments. SPA and Tomo more frequently exhibited longer "hard" and "soft missing" regions, with 74% and 88% in SPA, and 77% and 86% in Tomo, respectively. This reveals Cryo-EM’s greater acceptance of prolonged missing segments compared to X-ray.

Regarding compositional analysis of residues (S1 Fig), X-ray entries largely comprised "modeled" residues (93.5%), with minor proportions of "hard missing" (5.5%) and "soft missing" (1%) residues. SPA and Tomo had different distributions, featuring 75.9% "modeled" residues, 22.1% "hard missing", and 2% "soft missing" in SPA, and 77.5% "modeled", 20.4% "hard missing", and 2.1% "soft missing" in Tomo. The Median counts of "hard missing" residues per entry were 5 for X-ray, 25 for SPA, and 11 for Tomo, also illustrating X-ray’s less favor for disorder residues in models compared with Cryo-EM.

Overall, by this dataset it reveals notable variability in how experimental methods address structural disorder. SPA and Tomo offer greater tolerance for disordered regions compared to X-ray, managing longer unstructured segments and a higher proportion of missing residues.

### Correlation of Features with Residue Groups Using Pearson Analysis

Building on distribution analysis of "modeled" and "missing" residues, we conducted a correlation analysis to identify features that distinguish between these groups across different experimental methods. We examined residue counts alongside specific feature counts for each entry, conducting Pearson correlation tests to identify features strongly indicative of "modeled", "hard missing" and "soft missing" group memberships.

As in Fig 1, for "modeled" residues, the most correlated feature was the count of residues with high pLDDT scores (≥ 70), exhibiting strong correlations of 0.99, 0.92 and 0.90 for X-ray, SPA and Tomo, respectively. Additionally, the count of residues showing ordered IUPred scores (< 0.5) also demonstrated strong correlations of 0.98, 0.92, and 0.90.

**Fig 1.**
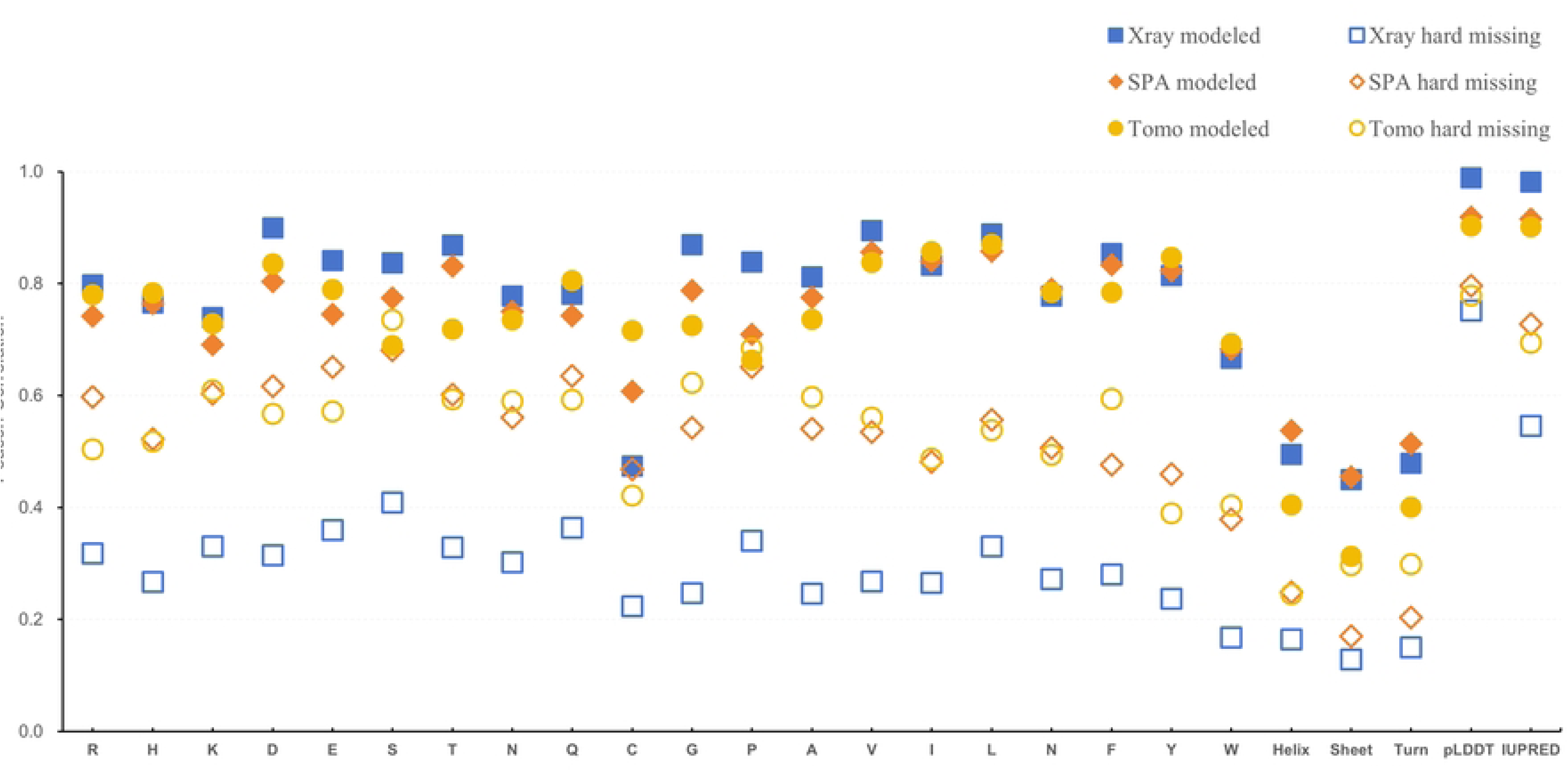
Correlation analysis of “modeled” and “hard missing” with structural features in X-ray, SPA and Tomo experiments. In "Modeled" residues (solid), the correlations observed with high pLDDT scores (≥ 70), displaying Pearson correlations of 0.99 (X-ray), 0.92 (SPA) and 0.90 (Tomo), and with ordered IUPred scores (< 0.5), showing correlations of 0.98 (X-ray), 0.92 (SPA), and 0.90 (Tomo). The next correlation groups are amino acid residue types, then followed by the residue counts of secondary structure types. In "Hard missing" residues (hollow), highest correlation is found with low pLDDT scores (< 70) and disordered IUPred scores (> 0.5), with correlations of 0.75 and 0.55 for X-ray, 0.80 and 0.73 for SPA, and 0.78 and 0.69 for Tomo. The amino acid residue types and counts of secondary structure types show slight correlations. The "Soft missing" residue (not showed) group show minimum correlations from 0.1-0.3 for all these features. Xray, square; SPA, diamond; Tomo, circle.

Similarly, "hard missing" residues correlated with counts of low pLDDT (< 70) and disordered IUPred scores (≥ 0.5), yielding correlations of 0.75 and 0.55 for X-ray, 0.80 and 0.73 for SPA, and 0.78 and 0.69 for Tomo. These results underscore the relationship between model defects and these predictive metrics.

Conversely, the "soft missing" group did not display obvious linear correlations with any feature tested across the experimental setups, suggesting that its characteristic complexity is not fully captured by straightforward correlation measures.

Beyond the highlighted pLDDT and IUPred scores, certain amino acid composition metrics showed correlations with the "modeled" group and a moderate association with "hard missing" group. However, these correlations remain negligible for "soft missing" group, indicating a broader diversity in this category. To be specific, amino acid compositions linked Serine and Alanine with high occupancy in “hard missing” group, while Tryptophan and Cysteine showed low presence across X-ray/SPA/Tomo. In contrast, “soft missing” residues varied more, with high presence of Glutamate, Lysine, and Alanine, and low presence of Tryptophan, Cysteine, and Methionine. In “soft missing” group, Serine and Alanine also appear frequently.

Notably, counts of residues forming secondary structure elements (such as helices, sheets, and turns) did not exhibit correlations comparably to other features, suggesting that measuring counts of secondary structure type alone is insufficient for predicting structural presence.

Overall, this analysis reveals that pLDDT and IUPred scores are useful for differentiating “modeled” from “hard missing” residues but are less effective for “soft missing” residues. Additionally, the composition of specific amino acids is a notable factor.

### Insights into the Landscape of pLDDT and IUPred Scores Across Residue Groups and Experiments

Based on correlation analysis found key features, pLDDTs and IUPRed scores were investigated by their distribution in different groups across X-ray, SPA, and Tomo datasets, to understand how these scores reflect prediction of structural reliability and disorder, providing overview for classifying "modeled" versus "missing" residues.

Residues were categorized into four quadrants based on pLDDT and IUPred scores: Q1 consists of residues with high pLDDT confidence and IUPred order; Q2 includes residues with high pLDDT confidence but IUPred disorder; Q3 comprises residues with low pLDDT confidence but IUPred order; and Q4 contains residues with low pLDDT confidence and IUPred disorder.

Fig 2 and Table S1 summarizes these findings. In X-ray data, "modeled" residues predominantly reside in Q1 (93.6%) with a minor presence in Q2. "Hard missing" residues are more evenly distributed to low confidence area, particularly in Q3, indicating structural uncertainty. "Soft missing" residues mostly appear in Q1 (63.7%), with some distribution in Q3 (20.1%).

**Fig 2.**
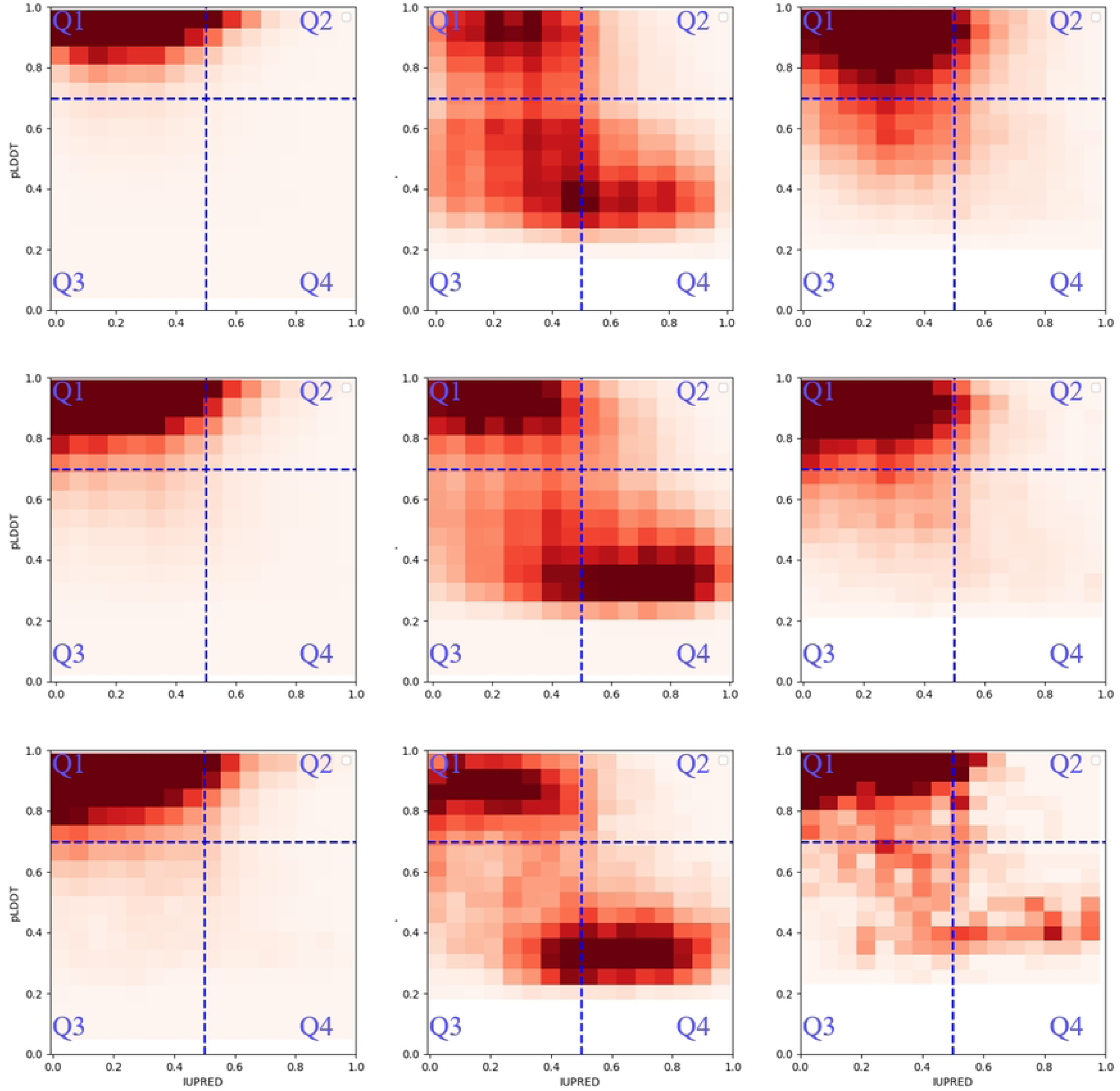
Distribution of residue categories across quartiles in X-ray, SPA, and Tomo Data. The distribution of "Modeled" (Left), "Hard missing" (Middle) and "Soft missing" (Right) residue group split with Q1 to Q4 across X-ray (Up row), SPA (Middle row), and Tomo (Down row) are represented as a density heatmap. Darker color corresponds for higher condensed population.

In SPA data, "modeled" residues cluster in Q1 (86.6%) with slight increases in Q2 and Q3 compared to X-ray data. "Hard missing" residues demonstrate a similar distribution to X-ray, spread across Q1, Q3, and Q4, albeit with varying intensity. "Soft missing" residues are primarily in Q1 (73.2%), with a reduced presence in Q3 and minimal representation in Q2 and Q4.

For Tomo data, "modeled" residues mainly occupy Q1 (84.2%), consistent with X-ray and SPA trends. "Hard missing" residues are distributed across Q1, Q3, and Q4, with a slightly reduced concentration in Q3. "Soft missing" residues are largely found in Q1 (65.3%), with notable representation in Q3 and increased presence in Q4 compared to X-ray and SPA datasets.

Overall, in “modeled” group, Q1 consistently houses the majority of residues in all methods, indicating that high pLDDT and low IUPred scores are strong indicators of structurally stable and ordered regions. The similarity in distribution patterns across different methods suggests that these techniques similarly shared predicted confidence and disorder scores.

The "hard missing" group shows more variability, particularly in Q3 and Q4, where lower pLDDT scores indicate uncertainty in structural predictions. The distribution across Q1, Q3, and Q4 underscores the challenges in predicting these residues. Furthermore, a decline in prediction confidence is observed in Q3, which does not correlate with the slope of disorder prediction metrics compared to Q2. This divergence suggests the potential existence of an underlying relationship between these two different prediction factors that warrants further investigation. "Soft missing" residues demonstrate intermediate patterns between the other two groups, clustered predominantly in Q1 but also appearing in Q3 across all methods. The Tomo data exhibits a higher presence of "soft missing" residues in Q4, suggesting increased disorder and low confidence in structure prediction compared to X-ray and SPA, possibly due to the differing progress used in tomographic and comparable lower resolution reconstructed structure.

These findings highlight the distinct characteristics of pLDDT and IUPred scores for "modeled" versus "missing" residues, illustrating how these scores are distributed among partially or fully structured residues and those found in inherently disordered or ambiguous regions.

### Predictive Modeling of Residue Groups from 2022 to 2023

Based on the analysis of residue features, another key question worth investigation is how the pLDDT and IUPred scores correlate with the missing coordinates across various structural experiments, especially following the release of the prediction algorithms. We evaluated the predictive capabilities using two fundamental models (pLDDT and IUPred) and an advanced LSTM model, focusing on data collected from 2022 to 2023.

Initially, a simple hypothesis was formulated based on prior analysis to classify residues: those with pLDDT scores below 70 or IUPred scores above 0.5 are considered "hard missing" while others are classified as "modeled". The "soft missing" group was temporally excluded from this classification due to their similarity to "modeled" residues, as indicated by previous heatmaps and lower correlation in this dataset.

To classify "soft missing" residues, we trained an LSTM model and incorporate sequence context. The dataset was split into a training set (entries before 2022) and a validation set (2022-2023 entries). The LSTM was trained on sequences along with pLDDT and IUPred scores, and then validated on the latest data.

Performance comparisons of these models on groups of residues were made using precision, recall, and F1 scores, summarized in Table 2. For "modeled" residues, the LSTM model excelled, achieving F1-scores of 0.98 (X-ray), 0.91 (SPA), and 0.91 (Tomo), closely followed by the pLDDT model. The IUPred model performed the least well.

**Table 2.**
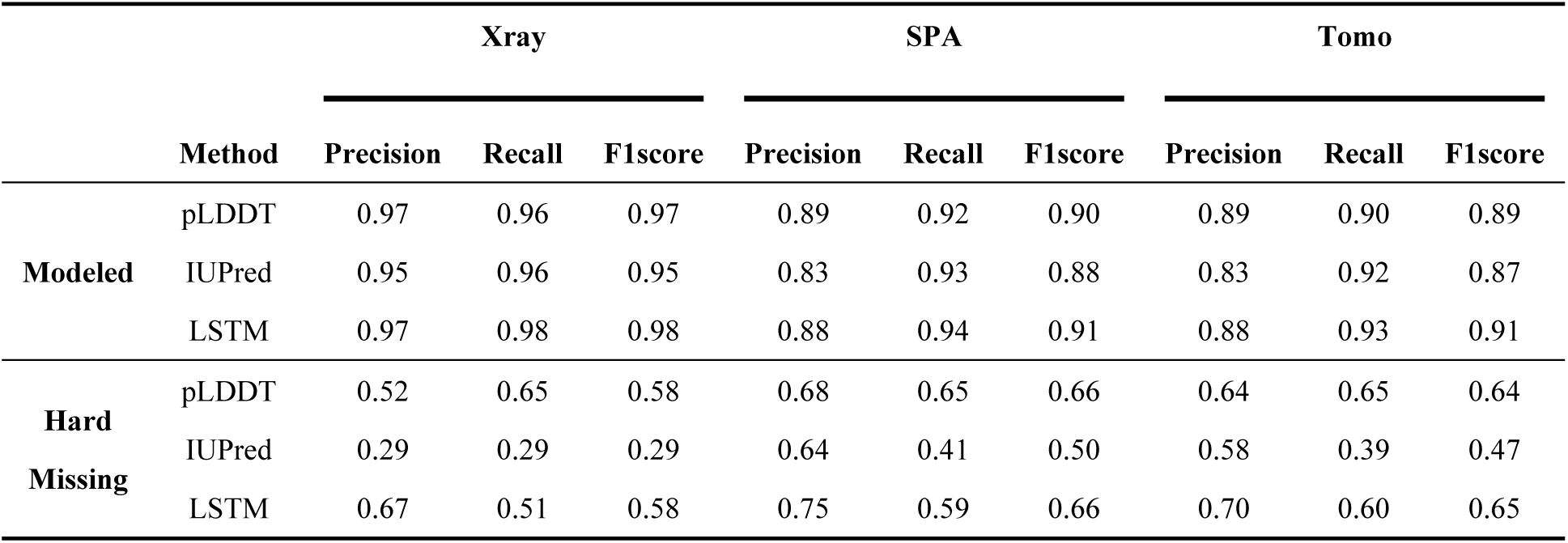
Summary of residue groups predicted by different methods.

|  |  | Xray |  |  | SPA |  |  | Tomo |  |  |
| --- | --- | --- | --- | --- | --- | --- | --- | --- | --- | --- |
|  | Method | Precision | Recall | F1score | Precision | Recall | F1score | Precision | Recall | F1score |
| **Modeled** | pLDDT | 0.97 | 0.96 | 0.97 | 0.89 | 0.92 | 0.90 | 0.89 | 0.90 | 0.89 |
|  | IUPred | 0.95 | 0.96 | 0.95 | 0.83 | 0.93 | 0.88 | 0.83 | 0.92 | 0.87 |
|  | LSTM | 0.97 | 0.98 | 0.98 | 0.88 | 0.94 | 0.91 | 0.88 | 0.93 | 0.91 |
| **Hard Missing** | pLDDT | 0.52 | 0.65 | 0.58 | 0.68 | 0.65 | 0.66 | 0.64 | 0.65 | 0.64 |
|  | IUPred | 0.29 | 0.29 | 0.29 | 0.64 | 0.41 | 0.50 | 0.58 | 0.39 | 0.47 |
|  | LSTM | 0.67 | 0.51 | 0.58 | 0.75 | 0.59 | 0.66 | 0.70 | 0.60 | 0.65 |

In predicting "hard missing" residues, the LSTM also led with F1-scores of 0.58 (X-ray) and 0.66 (SPA), marginally surpassing the pLDDT model for Tomo, with scores of 0.65 vs. 0.64. For "soft missing" residues, however, the LSTM did not classify any residues to this group during validation. This is possibly due to previously observed low correlation efficiency and scarce of data points.

Additionally, we evaluated prediction performance for "hard missing" regions, distinguishing between "short" and "long" categories in Table 3. Using a medium overlap (>50%) between true and predicted regions, the LSTM’s F1-scores for "short" regions were 0.75/0.82/0.79 (X-ray/SPA/Tomo), close to the pLDDT’s 0.75/0.78/0.80. For "long" regions, the pLDDT model held slight advantages with F1-scores of 0.65/0.76/0.70 over the LSTM’s 0.61/0.75/0.69.

**Table 3.**
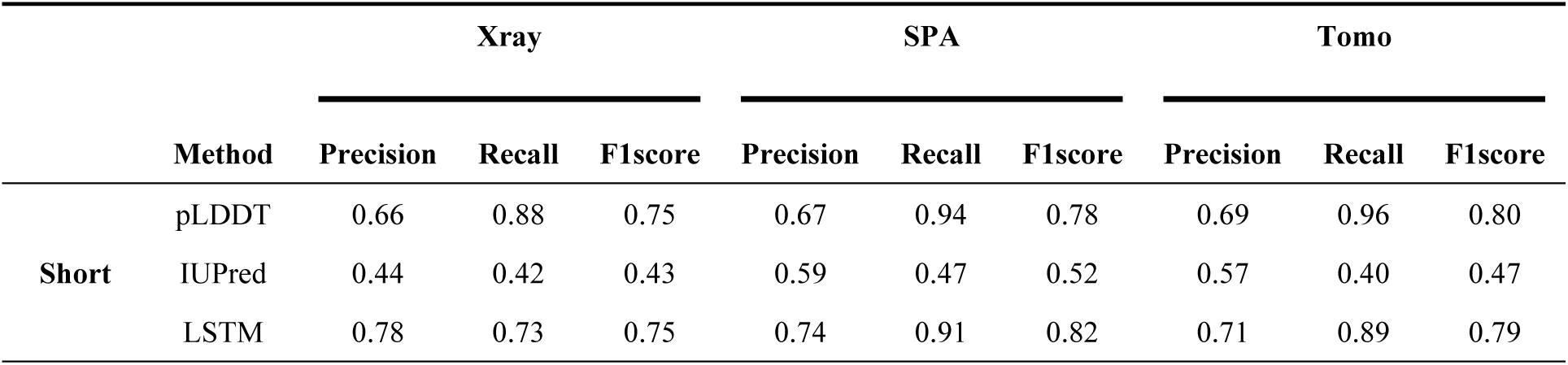
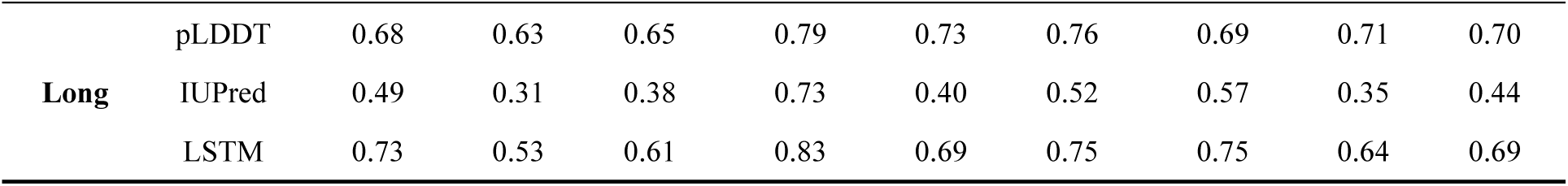
Summary of “Hard missing” regions predicted by different methods.

Both the basic pLDDT approach and the LSTM model have shown moderate performance in classifying “hard missing” residues, underscoring their capability in identifying likely unstructured regions with acceptable accuracy. The neural network model, in particular, holds considerable potential for enhancement. With the prospective integration of additional data sources and refinement of training methodologies, its promise as a sophisticated tool for this task is evident, poised for even greater performance gains in subsequent studies. On the other hand, the application of pLDDTs in predicting extended segments of protein structures provides a reliable baseline method.

Overall, the variation in performance across different groups emphasizes the differing correlations among models based on experimental methods, underscoring a comparably more robust correlation within SPA. By evaluating these findings, it is revealed that different approaches could be made to preview dependable and efficient routes to addressing the complexities inherent in modeling expansive and challenging regions of proteins.

### Application of pLDDT and LSTM Models in Analyzing Human Protein Dataset

The human protein SLC9A3 (Sodium/hydrogen exchanger 3, Uniprot P48764) is made as an example to illustrate the efficacy of different models in classifying "modeled" and "hard missing" residues. As depicted in Fig 3A, the structure includes spanning residues 40 to 665 (PDB: 7X2U), with "hard missing" regions between position 466-473 ("short") and 539-616 ("long"), highlighting overall agreement between predicted and experimental structural results. Intriguingly, both models identified region 616-665 as possibly missed in structures, which is actually a key functional region related to interaction, therefore reflecting dynamics and structural flexibility.

**Fig 3.**
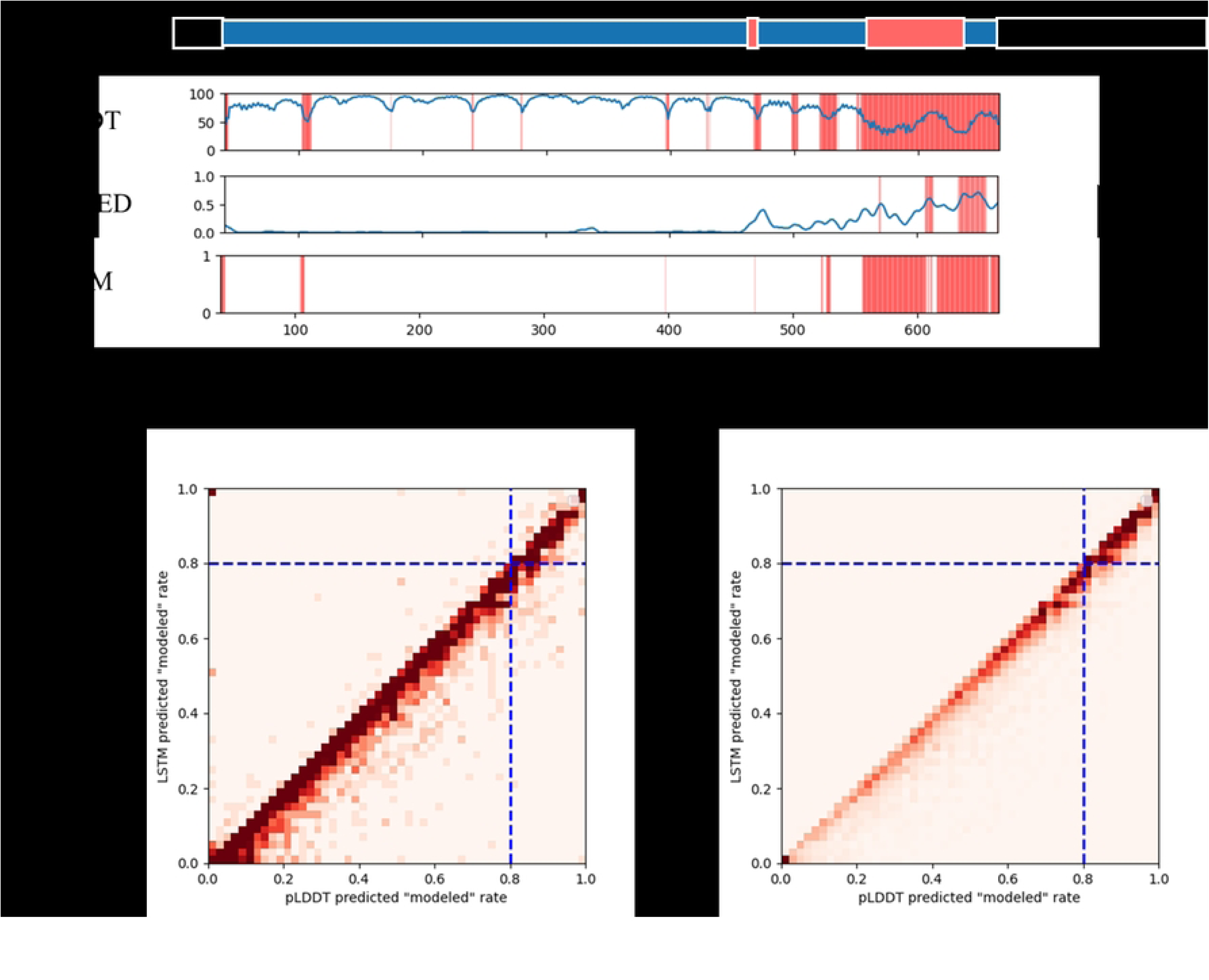
Example of prediction results and predicted “modeled” residue of human protein. (A) The full-length example of the protein SLC9A3 with structure solved (7X2U_1) and aligned with prediction results of models is depicted, as “modeled” (blue) segments, "hard missing" (red) and rest of unmodeled segments (black). (B) The density heatmap plots the predicted "modeled" rates from the pLDDT and LSTM models, on human protein dataset. The blue dashed lines indicate thresholds for high confidence in "modeled" ( ≥ 80%) predictions, including 322 "unsolved sequence" entries (left) and 3179 "unstructured sequence" entries exceeding 200 residues (right).

To expand application, it is targeted on unsolved human proteins, aiming to address the unrecorded ones while with high potential of be structured regions. Entries were curated into two categories: "unsolved sequence" comprising 8099 partially solved entries, and "unstructured sequence," containing 11665 entries without PDB representation until March 2024. The goal was to locate residues and regions in these unknown sequences with a high potential to be "modeled" to facilitate future structural studies. Using the basic pLDDT model and trained LSTM model, we assessed unknown regions, prioritizing those predicted to be "modeled" ratio, as detailed in Fig 3B. As showed, the comparison of results between pLDDT and LSTM demonstrated a strong correlation in their predicted "modeled" rates, suggesting that the trained LSTM effectively calibrates pLDDT predictions concerning the classification of missing residues.

Our findings revealed 3179 "unstructured sequence" entries with high potential for "modeled" residues (≥80% prediction by both models) with counts exceeding 200 residues, and 322 "unsolved sequence" entries showing a significant capacity for abundant "modeled" residues. This identifies a promising repository for future structural and functional experiments.

## Discussion

Disordered regions, while occasionally absent from structural maps, are known to indicate important protein function and present challenges in structural determination. Advances in structure predictions, notably through AlphaFold2, demonstrate a promising correlation between low-confidence regions and various disorder across diverse species [25, 31, 33, 34]. Evidence from AlphaFoldDB also supports the prevalence of low pLDDT scores in IDRs, highlighting the potential of using structure prediction methods to explore the largely unknown protein universe. This approach offers valuable insights into disordered regions related to human proteins, diseases, and the broader protein landscape [35–37].

Our investigation seeks to bridge the deficient knowledge identified in previous studies, which were often constrained by the limited prediction structural datasets available at the time or relied on earlier versions of disorder databases like DisProt [29, 31, 38]. By harnessing a dataset combining both recent and historical structural data produced through a range of key structural techniques, we thoroughly examine AlphaFold2’s predictive capabilities alongside other methods, including IUPred and LSTM. This comprehensive approach allows us to scrutinize the accuracy of these methods, particularly in their ability to elucidate intricate protein segments that have consistently posed challenges to structural characterization. Our analysis not only enhances our grasp of these elusive regions but also provides a valuable cross-sectional view of prediction methodologies applied to a diverse array of experimental data, spanning from the historical to the contemporary.

By classifying residues into “modeled”, “hard missing” and “soft missing”, we shed light on the intricate relationships between types of missing residues, regions, and the distribution of prediction scores. “Modeled” residues, which generally achieve high pLDDT scores, are found in areas where AlphaFold2’s predictions are most reliable, particularly in data derived from X-ray crystallography, which exhibits the highest median pLDDT scores. On the other hand, “hard missing” residues largely fall within low-confidence regions, underscoring their inherent unpredictability for structure prediction models. The distributions of IUPred scores further helps draw a clear distinction between predicted ordered and disordered regions, indicating different predictability levels behind. “Soft missing” residues represent a boundary case, potentially oscillating between order and disorder, complicating predictions due to their dynamic nature and potential roles in the assembly of protein complexes under specific environmental conditions [29, 39]. It’s important to note that, within our extensive dataset focusing on historically challenging hard-to-structure residues and regions, “soft missing” residues are presumed to include many that are borderline adjacent to hard-to-structure regions, while “hard missing” ones are situated within these difficult regions in this study.

While our study did not identify a direct link between the counts of residues within each secondary structure and the occurrence of missing residues, AlphaFold2 has demonstrated effectiveness in predicting secondary structures such as α-helices and β-sheets. It has been observed that residues within different secondary structures tend to exhibit varied pLDDT distributions, indicating the necessity for a more precise classification of regions, and dynamic analysis complemented by experimental validation techniques to enhance our understanding of these complex structural behaviors [25, 40].

Re-evaluating feature such as pLDDTs distributions across experimental methods reveals structural determination challenges tied to specific techniques. For example, while X-ray crystallography excels in resolved regions with high-pLDDT, cryo-EM mitigates crystallization challenges, highlighting its utility in more diverse experimental conditions. The reason behind might be, high pLDDT scores are generally associated with high confidence in the predicted structures and may indicate regions that are folded. Conversely, low pLDDT scores often point to regions of increased flexibility and heterogeneity on conditions, which present a challenge for precise structural prediction due to additional factors needed to be considered such as the related homologous sequences or composition bias [34, 38, 39].

Intriguingly, within our dataset, we observe fewer instances where disorder predicted residues have high structural prediction confidence than instances where residues are predicted to be ordered with low prediction confidence. This pattern spans across various groups of residues and structural experimental outcomes, suggesting that in these cases, a prediction of disorder might be more consistently aligned with low structural prediction confidence, while the reverse scenario. Such findings highlight the importance of continual updates and re-evaluations of disorder annotations, as mentioned by newly released DisProt [41].

Incorporating sequence characteristics alongside pLDDT/IUPred scores into neural network models has refined the ability to predict protein structures. By integrating more features and optimized network, this strategy may further enhance the identification of elusive targets, particularly within disorderly proteins where conventional approaches falter. Understanding the correlations between predicted and experimental structures, especially after the introduction of AlphaFold, enables us to better comprehend and pinpoint structural targets in proteins yet to be solved. This insight allows for more reasonable designing of experiments, directing future research opportunities towards areas of proteins with high scores indicative of stability, or low scores signaling unstructured regions but with potential functional importance.

While AlphaFold provides significant predicted structural insights, exclusive reliance on prediction has risks in certain cases [27, 39]. To obtain a more complete picture of protein behaviors, it is imperative to integrate structure predictions with other information, such as experimental validations from dynamic measuring methodologies, which can provide valuable insights to understand these “unstructured” in protein with flexibility and complex assemble, and other environmental factors which have significant influences on structures and functions.

## Conclusions

This study highlights methodological advancements in missing coordinates correlation analysis, particularly for hard-to-structure related "hard missing" regions, using AlphaFold2’s pLDDT scores in conjunction with advanced modeling techniques. This work provides a robust framework for systematic analysis and preview of missing residues and regions in protein structures, illustrating the intricate relationship between structural determination methods and disorder by correlated prediction. Integrating pLDDT and LSTM models aligns with experimental data while identifying areas in protein dynamics needing refinement, especially in novel ones. This research enhances predictive strategies crucial for exploring unresolved protein regions, and setting the stage for improved experimental designs and insights into protein dynamics relevant to both computational and experimental research.

## Acknowledgments

I would like to express my sincere gratitude to Dr. Zhanyu Ding from Shanghai YueXin Life-Science Information Technology Co. Ltd. for her generous support in providing resources and guidance for the experimental data analysis in this study. I also extend my appreciation to my colleagues in the cryo-electron microscopy team for their valuable suggestions

## Supporting information

**S1 Fig. Compositional Distribution of Residue Categories in Structural Determination Methods.** The figure displays the counts of residues classified as "Modeled" (**left**), "Hard missing" (**middle**), and "Soft missing" (**right**) across various structural experiments, including X-ray (**blue**), SPA (**orange**), and Tomo (**yellow**). The median counts of "Modeled" residues per protein are 242 for X-ray, 190 for SPA, and 179 for Tomo. For "Hard missing" residues, the median counts are 5 for X-ray, 25 for SPA, and 11 for Tomo. Median counts for "Soft missing" residues are not reported.

**S2 Fig. Distribution of Residue Scores in Different Experimental Methods.**

A. The distribution of pLDDT residue scores (ranging from 0 to 100) across the "Modeled," "Hard missing," and "Soft missing" groups is presented for X-ray, Single-Particle Analysis (SPA), and Tomography (Tomo) experiments. In the X-ray dataset, the "Modeled" group shows the highest median pLDDT score at 97.1, followed by the "Soft missing" group at 84.4 and the "Hard missing" group at 55.5. For the SPA dataset, the medians are 92.7 for "Modeled," 86.7 for "Soft missing," and 54.5 for "Hard missing." In the Tomo dataset, medians are 91.5 for "Modeled," 88.6 for "Soft missing," and 52.3 for "Hard missing."
B. The distribution of IUPred scores (ranging from 0 to 1, with scores over 0.5 indicating disorder) is shown for the same groups and experimental methods. In the X-ray experiment, the "Modeled" group has the lowest median IUPred score at 0.2, followed by "Soft missing" at 0.29 and "Hard missing" at 0.38. In SPA, the median IUPred scores are 0.19 for "Modeled," 0.23 for "Soft missing," and 0.43 for "Hard missing." In Tomo, the scores are 0.2 for "Modeled," 0.32 for "Soft missing," and 0.42 for "Hard missing." These findings highlight distinct score distributions across experimental methods and residue categories.

**S1 Table. Distribution of residues classified by types from different methods**

